# Demonstration of Bioplastic Production from CO_2_ and Formate using the Reductive Glycine Pathway in *E. coli*

**DOI:** 10.1101/2023.12.02.569694

**Authors:** Daria Fedorova, Roee Ben-Nissan, Eliya Milshtein, Ghil Jona, Nili Dezorella, Gil Feiguelman, Rostislav Fedorov, Aya Gomaa, Ariel B. Lindner, Elad Noor, Ron Milo

## Abstract

There is a strong need to develop technologies that reduce anthropogenic pollution and the dependence on nonrenewable Earth resources. One way of doing so is by harnessing biological systems for replacing the production of fossil-fuel based goods with low-environmental-impact alternatives. Recently, progress was made in engineering the model organism *E. coli* to grow using CO_2_ and formate as its only carbon and energy sources using the reductive glycine pathway (rGlyP). Here, we use this engineered strain of *E. coli* as a host system for the production of polyhydroxybutyrate (PHB), a biologically derived and biodegradable plastic. We confirmed the production of PHB in this strain using Nile red fluorescent microscopy, transmission electron microscopy, and GC measurements. Since formate can be efficiently generated from CO_2_ by electrochemical reduction using renewable energy sources, this study serves as a proof of concept for the emerging field of electro-bioproduction.

## Introduction

Currently, a large fraction of essential chemicals are produced from natural gas and petroleum. These petrochemicals (such as ethylene and acetylene) are used as a feedstock for the production of plastics, fuel, textiles, cosmetics and fertilizers (International Energy Agency 2018). Using petroleum as a feedstock poses major challenges, both because it is a finite resource that is becoming increasingly expensive to extract, and because petrochemicals are responsible for 2% of global greenhouse gas emissions (Ritchie and Roser 2023).

Alternatives to petroleum, based on renewable feedstocks such as maize, sugar cane, or other plant-based sources, have been gaining momentum in recent years. However, these bioproducts have a large land and water footprint and often compete with food production. This could have consequences for food security, especially in the context of a growing population and projected yield reductions from climate change (Hultgren et al. 2022). For example, if we were to replace the petrochemical-based plastics used for packaging purposes (which account for 44% of global plastic consumption) with bioplastics derived from plant material, it would require a minimum of 61 million hectares of land, adding more than 5% to the total cropland area currently used worldwide (Brizga, Hubacek, and Feng 2020). Expanding cropland has a negative effect on biodiversity due to loss and fragmentation of natural habitats, and has to be avoided (Zabel et al. 2019; Foley et al. 2005; Chaplin-Kramer et al. 2015). There is therefore a need to find a different alternative to bioproduction of chemicals which is scalable, efficient, and decoupled from human food production.

### Poly-β-hydroxybutyrate (PHB) production in microorganisms

Poly-β-hydroxybutyrate (PHB) is a naturally biodegradable polyester, synthesized natively in various microorganisms as a store of energy and carbon under nutrient-depleted conditions. Humanity mainly uses PHB for food packaging, as well as biocompatible and biodegradable implants, medical scaffolds, and encapsulation of medicines (dos Santos et al. 2017; Nagarajan et al. 2021). PHB can be produced in *E. coli* heterologously by expression of a PHB operon. As shown in Figure 1, PHB production in *E. coli* requires expression of three non-native enzymes: PhaA (3-ketothiolase), PhaB (acetoacetyl-CoA reductase) and PhaC (PHB synthase). PhaA condenses two molecules of acetyl-CoA to form acetoacetyl-CoA, which is then reduced by PhaB to create (R)-3-hydroxybutyl-CoA (3HB). PhaC then uses 3HB as the monomer for polymerising poly-3-hydroxybutyrate (PHB). On an industrial scale, PHB is produced heterotrophically in fermenters by bacteria such as *Cupriavidus necator* or recombinant *Escherichia coli*, using glucose as a feedstock (G.-Q. Chen 2009). *C. necator* is also capable of producing PHB natively from CO_2_ as the sole carbon source (using aerobic respiration of H_2_ as the energy source) (Soyoung Kim et al. 2022) (Garcia-Gonzalez et al. 2015). However, unlike *E. coli, C. necator* has a poorly characterized genome and limited genome manipulation tools. This hinders the use of *C. necator* for producing non-native chemicals. Therefore, we aimed to demonstrate a synthetically engineered C1-feeding *E. coli* as a platform for PHB bioproduction. This serves as a proof-of-concept for bioproduction of other biochemicals, assuming their pathways can be readily introduced into this well-characterized model organism.

**Figure 1:**
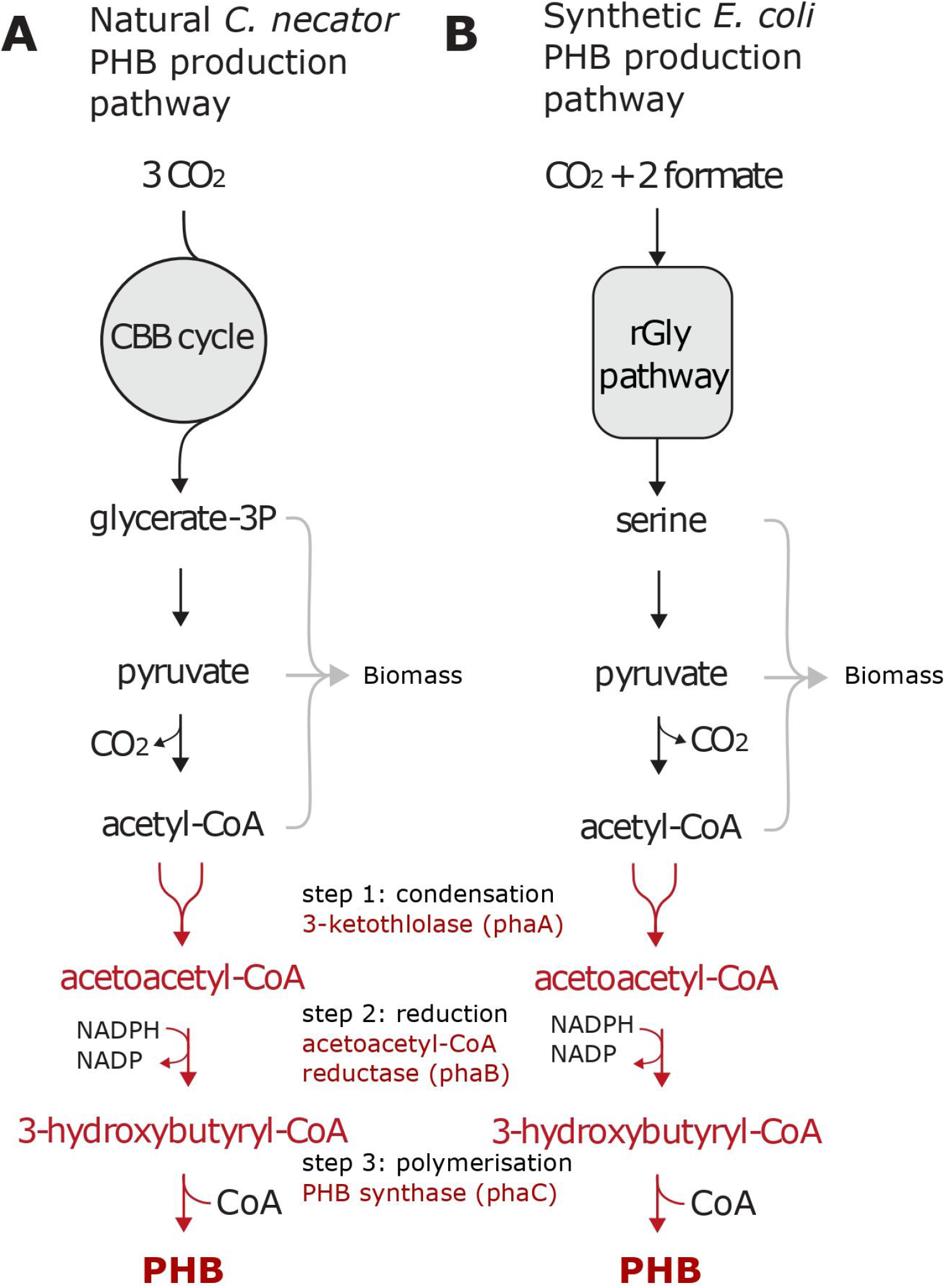
Metabolic pathway of PHB production in *C. necator* and *rGlyP E. coli*. **(A)** Metabolic pathway of PHB production in C. necator using the Calvin-Benson-Bassham (CBB) cycle for carbon assimilation. The PHB pathway is highlighted in red. **(B)** Metabolic design of rGlyP E. coli producing PHB. Formate and CO_2_ are assimilated into carbon metabolism using the reductive Glycine pathway. PHB is produced from Acetyl-CoA by heterologous expression of the PHB operon (in red).

### Engineering E. coli to feed on C1 compounds

Recently, progress has been made in engineering *E. coli* to feed on C1 compounds (Seohyoung Kim et al. 2020; Gleizer et al. 2019; F. Y.-H. Chen et al. 2020; Keller et al. 2022). As a background to our work using one such *E. coli* strain, we highlight three studies that demonstrated growth on C1 feedstocks using the reductive glycine pathway (rGlyP), the reductive pentose phosphate (rPP) cycle, and the ribulose monophosphate (RuMP) cycle.

(Seohyoung Kim et al. 2020) introduced the reductive glycine pathway (rGlyP), an efficient route for direct formate and CO_2_ assimilation into central metabolism (Fig. 1, left). By combining rGlyP with laboratory evolution approaches, they successfully achieved formatotrophic growth of synthetically engineered *E. coli*. This strain had a doubling time of less than 8 hours and a biomass yield of 2.3 gCDW per mol of formate. This growth is strongly coupled to formate as alternative electron donors are not effective with rGlyP. Additionally, this strain lacks a CO_2_ concentrating mechanism, requiring artificially high concentrations of CO_2_ for biomass accumulation.

(Gleizer et al. 2019) established complete synthetic autotrophy in *E. coli* by combining rational design and adaptive laboratory evolution. The evolved strain uses the reductive pentose phosphate cycle (rPP) cycle for all its biomass production and relies solely on CO_2_ and formate as its carbon and energy sources respectively. This strain displays a doubling time of 18 ± 4 hours and has a formate-to-biomass conversion yield of 2.8 ± 0.8 gCDW/mol formate. Further engineering of this strain may enable growth at ambient CO_2_ levels, which is useful in industrial settings. Furthermore, since carbon fixation was decoupled from the energy supply in this strain, the system is modular and can be readily adapted for the use of other electron donors (*e.g*., methanol, hydrogen).

In another study, (Keller et al. 2022) transformed *E. coli* into a synthetic methylotroph that assimilates methanol via the energy-efficient ribulose monophosphate (RuMP) cycle. Methylotrophy was achieved after evolving a methanol-dependent *E. coli* strain over 250 generations in a continuous chemostat culture, using methanol as the sole source of carbon and energy. This strain is strongly coupled to methanol and cannot use any other energy source. Recently, this strain was engineered to demonstrate the production of four diverse biochemicals, one of which is PHB (Reiter et al. 2024).

In light of the studies described above, we opted to incorporate a PHB production pathway from *C. necator* into the *E. coli* strain that was previously equipped with rGlyP (Seohyoung Kim et al. 2020). While the strain that utilizes the RuMP cycle for growth on methanol supports relatively rapid growth, our aim was to showcase the production of PHB from formate, as it corresponds with the concept of a formate economy (as elaborated in the discussion section). Although both the autotrophic strain and the rGlyP strain are suitable for this purpose, the latter exhibited a significantly faster growth rate and is more amenable to straightforward genetic engineering

## Results

*C. necator* H16 naturally produces polyhydroxybutyrate (PHB) from acetyl-CoA as a carbon storage polymer that accumulates as granules in the cytoplasm. The three genes required for PHB production are located within a single operon. We set out to introduce the PHB production pathway into formate-utilizing strains of *E. coli* that rely solely on formate and CO_2_ as their sources of carbon and energy by using the reductive glycine pathway (rGlyP). rGlyP is a synthetic pathway that offers an efficient route for the direct assimilation of formate and CO_2_ into the rest of cellular metabolism (Fig. 1B). Notably, a variant of this pathway was identified in the anaerobic bacterium *Desulfovibrio desulfuricans* (Sánchez-Andrea et al. 2020). Building upon previous work by (Seohyoung Kim et al. 2020), who successfully achieved formatotrophic growth in the engineered *E. coli*, we sought to adapt this strain for production of PHB. We validated that this strain grows quickly on rich media, making it amenable to fast genetic manipulation. Furthermore, its doubling time on M9 minimal media, supplemented with formate and CO_2_ as the only reducing and carbon sources, was less than 8 hours, sufficiently fast for our experimental setup. To produce PHB in *E. coli*, we introduced a pHB-4 plasmid (Liu et al. 2020) containing the PHB operon under the control of the auto-inducible promoter PthrC3 (Liu et al. 2020). This promoter was selected due to its advantages over traditional inducible promoters, such as pT7 (Anilionyte et al. 2018). PthrC3 is an endogenous, short *E. coli* promoter that can auto-induce during early exponential growth, thereby enhancing efficiency and cost-effectiveness by eliminating the need for inducer chemicals in the media, which could also potentially be used as a carbon source (e.g., in Arabinose-inducible systems).

The formatotrophic strain carrying the PHB genes was cultivated in a fermenter using minimal media supplemented with 60 mM formate (as was previously determined to be the optimal concentration (Seohyoung Kim et al. 2020)) and an overlay of 10% CO_2_ in the headspace as the sole carbon source. The PHB-producing strain cultivated in the fermenter exhibited a prolonged lag phase compared to the non-PHB producing strain, suggesting that the synthesis of PHB slows bacterial growth (Fig. 2B). This was expected, as it is well-established that heterologous production of chemicals or proteins tends to slow microbial growth (Glick 1995).

**Figure 2:**
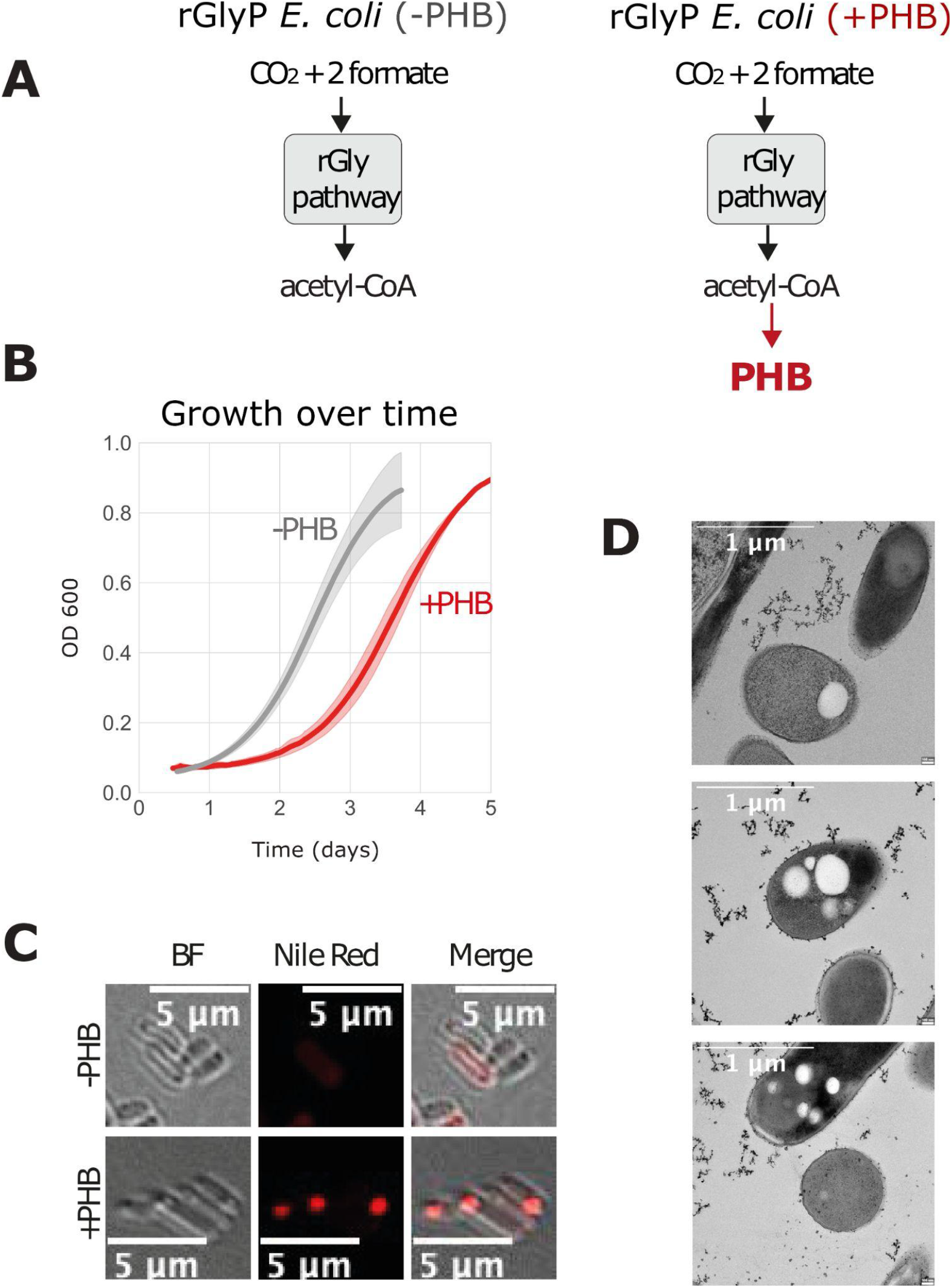
Validation of bioplastic production in *rGlyP E. coli*. **(A)** Metabolic design of rGlyP E. coli with and without PHB pathway. **(B)** Growth of isolated PHB producing strain in liquid M9 minimal media with 60 mM sodium formate and bubbled with a gas mixture of 10% CO_2_, 90% air (in duplicates). Growth was carried out in DASGIP fermenters (150 mL working volume). **(C)** Production of PHB in Formatotrophic E. coli as seen by confocal Nile red microscopy (red granules stained with Nile Red). Images were taken using a Nikon A1R HD25 confocal microscope, 60x. Images were cropped to show relevant fields of view and no post-processing was performed. **(D)** Transmission electron micrographs showing PHB granules inside the PHB producing rGlyP E. coli strain.

To confirm PHB production, we first employed the widely-used staining technique using Nile red (Juengert, Bresan, and Jendrossek 2018) (Kavitha et al. 2016) (Jendrossek, Selchow, and Hoppert 2007). Nile red is a stain that becomes fluorescent within hydrophobic cell structures, enabling detection through fluorescent confocal microscopy. Using this, we observed the formation of granules (in red) in a transformed strain of *E. coli* (Fig. 2C). In contrast, we did not observe such granules in a strain lacking the PHB-producing plasmid. As Nile Red also fluoresces when binding to bacterial membranes, we complemented these results with transmission electron microscopy, which reveals the morphology of the granules (Fig. 2D). Furthermore, to validate the chemical composition of granules is indeed PHB, we conducted Gas Chromatography with Flame Ionization Detection (GC-FID) analysis. Gas Chromatography is often used for the detection and quantification of PHB (Juengert, Bresan, and Jendrossek 2018) (Oehmen et al. 2005). We used it to verify the chemical structure of PHB using conventional PHB standards (Fig. S1), but we were unable to quantify the yield of PHB using GC-FID due to its low levels of accumulation (See Methods: Quantification of PHB by GC-FID).

## Discussion

It was previously demonstrated that *E. coli* can be engineered to grow on C1 compounds such as CO_2_, formate or methanol (Seohyoung Kim et al. 2020; Gleizer et al. 2019) (Keller et al. 2022). In this study, we use a synthetic formate-utilizing strain as a platform for the production of PHB. Through heterologous expression of the PHB producing pathway from *C. necator*, we successfully produced bio-plastic and confirmed its presence using analytical techniques.

The gold standard in the field of C1 metabolism is to perform 13C labeling experiments to ensure that all the carbon is derived from CO_2_ and formate (Gleizer et al. 2019; Seohyoung Kim et al. 2020). This method has already been used to demonstrate that the formatotrophic strain of *E. coli* synthesizes all of its biomass carbon exclusively from CO_2_ and formate in minimal media conditions. Since we used the identical strain described by Kim et al. 2020, under identical growth conditions (in both the growth and in the production phases) we assumed that all the carbon used for PHB production came exclusively from CO_2_ and formate as well, as these are the only sources of carbon available in the growth medium.

### Decoupling and other strategies for improving the production capacities

A high PHB yield is critical for financially competing with traditional plastic production as this greatly influences the cost of production. Here we examine various strategies aimed at maximizing PHB yields, thereby advancing its potential for commercial viability. We also note that a pipeline for rapid quantification of PHB will be instrumental for progress in these efforts.

As expected, the production of PHB is a burden on cell growth. It is known that production of value-added chemicals also leads to the accumulation of toxins and mutations (Borkowski et al. 2016). Expressing genes continuously requires a substantial amount of energy. It is more energy-efficient to express genes only when they are needed. To overcome these problems in bioproduction and reduce metabolic stress, it is common to decouple biomass accumulation from chemical production (Pouzet et al. 2020). Current decoupling strategies include using inducible promoters triggered by physical (e.g. temperature) (Menart et al. 2003) or chemical (e.g. pH, nutrient depletion, IPTG and galactose) signals (Chou et al. 1995) . Recently, complex circuit designs for dynamic regulation, such as optogenetics (Lalwani et al. 2021) and biosensing (Lo et al. 2016), were shown to successfully increase the product yield in several bioproduction strains. An additional decoupling strategy for synthetic or natively autotrophic organisms is to use a two-stage cultivation system (Koller 2018; Baumschabl et al. 2022) (Garcia-Gonzalez et al. 2015). This consists of a heterotrophic growth phase to reach a high OD and an autotrophic growth phase for production. For formatotrophs, different strategies can be used to decouple growth from production including using formate inducible promoters (Hanko et al. 2020). A two-stage cultivation system is a more robust and reliable approach for sustainable bioproduction compared to constitutive production of a molecule of interest. Further investigation is warranted to explore the potential of a two-stage cultivation system using synthetic C-1 feeding organisms.

Another approach to enhance PHB accumulation is by increasing the growth rate of the initial strains as it minimizes the fermentation costs and improves the overall robustness of the process. This can be accomplished through laboratory evolution or genetic modifications. Another viable strategy is metabolic engineering, involving the modification of bacterial metabolic pathways to boost PHB production. Specific genetic manipulations include modulating phasin expression, which is a surface-binding protein of polyhydroxyalkanoate (PHA) granules that is encoded by the phaP gene. Indeed, an E. coli strain harboring novel phasins from a highly productive PHB bacterium, *Halomonas sp*. YLGW01, exhibited increased PHB production by around 3-fold. (Lee et al. 2023).

Additionally, the deletion of *ldhA, pta*, and *adhE* genes, which encode enzymes in competing metabolic pathways, can be employed to redirect resources toward PHB production. The composition of the growth medium can exert a significant impact on PHB accumulation. By regulating factors like nitrogen and phosphorus concentrations, it is possible to create stress conditions that prompt bacteria to store excess carbon as PHB. Restricting these essential nutrients forces the cells to prioritize PHB production (Wang et al. 2009).

Modifying the physical characteristics of bacterial cells can also positively influence PHB accumulation. Strategies may encompass altering cell size and shape, division patterns or cell wall structures (Wu et al. 2016) . For instance, by overexpressing *sulA*, a cell division inhibitor, in *E. coli* harboring PHB synthesis operon, engineered cells became long filaments and accumulated more PHB compared with the wild-type (Wu, Chen, and Chen 2016).

Finally, *In vitro* polymerization of PHB could be another way for overcoming spatial constraints within the cytoplasm. Traditional PHB production occurs inside bacterial cells, however, developing techniques for *in vitro* (outside the cell) polymerization of PHB has the potential to streamline the production process. This approach may offer greater control over the polymer’s characteristics and reduce energy and resource consumption.

### Potential for photovoltaic-driven bioproduction

A cost-effective bioproduction method could utilize microorganisms that grow using molecules containing only a single carbon atom (C1), such as carbon dioxide, formate or methane. Recently, (Leger et al. 2021) analyzed the efficiency of using solar energy for converting CO_2_ derived from direct air capture into formate and other electron donors and using it as a feedstock for microbial production of protein for human consumption. The study found that photovoltaic-driven production of protein outperforms agricultural cultivation of staple crops in terms of caloric and protein yields per land area and could thus be used in the future to save limited land resources.

Electron carriers generated through electrolysis, such as formate and hydrogen, are essential in photovoltaic-driven production to shuttle electrons to microorganisms that grow on C1 feedstocks.

Formate can be produced by electrochemical reduction of CO_2_ with a Faradaic efficiency greater than 90% (Pletcher 2015; Taheri and Berben 2016). Compared to hydrogen, the electrochemical production of formate is less efficient, but it has the advantage of being soluble and non-flammable, and thus more convenient for large-scale usage.

In the electrochemical reduction of CO_2_, only carbon monoxide and formate exhibit efficient production (Jhong, Ma, and Kenis 2013) (Yishai et al. 2016). Since both require only a two-electron reduction of CO_2_, finding a suitable catalyst for their production is less challenging compared to multi-electron products such as methane, methanol, ethylene, oxalate, and acetate (Yoo et al. 2016) (Yishai et al. 2016). Exploring the bioproduction capabilities of organisms that use formate and CO_2_ as an energy and carbon source might therefore be a solution for achieving sustainable production of goods such as plastics.

Engineering an *E. coli* strain with rGlyP to heterologously express the PHB operon is a step in the long path towards electro-fermentation. To the best of our knowledge, this is the first report of bioproduction in a synthetic *E. coli* strain feeding on formate and CO_2_. This study serves as a proof of concept for synthetically engineering *E. coli* strains that can utilize formate and CO_2_ as a feedstock for the production of plastics and more generally the potential of photovoltaic-driven bioproduction for future sustainable production.

## Materials and methods

### Chemicals and reagents

Primers were synthesized by Sigma-Aldrich. PCR reactions were performed using KAPA HiFi HotStart ReadyMix or Taq Ready Mix. Glycine, Sodium formate, D-xylose, D-glucose, Nile-Red, PHB and 3HB standards were purchased from Sigma-Aldrich.

### Bacterial strains

The synthetic formatotrophic rGlyP *E.coli* strain was described previously by (Seohyoung Kim et al. 2020) and used as host organism for bioproduction of plastic by transformation of this strain with recombinant plasmid/s.

### Recombinant plasmids

pHB-4 was a gift from Kang Zhou (Addgene plasmid # 140957 ; http://n2t.net/addgene:140957 ; RRID:Addgene_140957)

### Fermentation of E. coli strains

The growth of the various strains was done in a DASBox multi-parallel fermentation system (Eppendorf). Briefly, 10 ml starters from each strain were used to inoculate each bioreactor that contained 140 ml minimal M9 media supplemented with vitamin B1, trace elements, and 60mM filter sterilized sodium formate, to an OD 600 of ∼0.05. Cultures were grown at 37 C with constant agitation and were supplemented with gas composed of 10% CO_2_ and 90% air. The growth of the culture was constantly monitored along the when the cultured reached saturation the cultures were diluted to the initial OD by aspirating the appropriate volume of the culture and replacing it with fresh M9 media as above. Samples from each strain were routinely collected.

### Growth tests of rGlyP E. coli strains

Method was adapted from (Seohyoung Kim et al. 2020)

### Sample preparation for Nile Red confocal microscopy

Nile red is a fluorescent dye used to visualize lipid-like inclusions. This dye binds to PHB granules and can be detected by fluorescence microscopy. Nile red staining was performed according to the protocol detailed in (Juengert, Bresan, and Jendrossek 2018). Cells were harvested by centrifugation (6,000x). 4 μl of cells were taken and pipet mixed with 1 μl of Nile red (10 μg/ml in DMSO). Then, 1 μl of the stained cell suspension was put on a microscopic slide and covered with an agarose pad. The cells were imaged using a Nikon A1R HD25 confocal microscope. Nile Red fluorescence was detected with DU4 detector, excitation of 561.5 nm laser, and emission filter 593/46 nm. Fluorescence intensity was adjusted to the negative control and imaged with the SR Plan Apo IR AC 60x water immersion objective (numerical aperture 1.27). Images were cropped to show relevant fields of view.

### PHB extraction, depolymerization and derivatization to methyl 3-hydroxybutanoate

Cell pellets were dried by lyophilization overnight, weighed, and transferred to an air-tight glass tube. 2 mL of 3% H_2_SO_4_ in methanol containing 190 µg/mL of benzoic acid as internal standard and 2 mL of chloroform were added to the pellet. PHB standards were prepared in chloroform and mixed 1:1 (v/v) with 3% H_2_SO_4_ in methanol containing benzoic acid. For PHB extraction, depolymerization by methanolysis, and derivatization, samples were incubated for 2.5 h in a boiling water bath. To initiate phase separation, 1 mL of ultra-pure water (MilliQ) was added to the solution and the samples subsequently incubated for 10 min in a sonication bath. The aqueous phase (upper) was discarded and the organic phase (lower) used for GC analysis.

### Quantification of PHB by GC-FID

By acidic methanolysis processed PHB (3-hydroxybutyrate methyl ester) was analyzed by gas chromatography (GC 6850, Agilent Technologies, Basel, Switzerland) equipped with a 7683B Series injector coupled to a flame ionization detector (FID). A DB-WAX column (15 m x 0.32 mm x 0.50 μm; Agilent Technologies, Basel, Switzerland) was used for metabolite separation with helium as the carrier gas at a flow of 30 mL/min. The following temperature gradient was applied: 3.5 min from 90 °C to 230 °C, 3 min at 230 °C, 1.25 min to 90 °C, 2.5 min at 90 °C. 1 uL of sample was injected. The split ratio was 2.0 and the detector temperature set to 270 °C. Peaks were confirmed by standards. Methyl benzoate was used as an internal standard to correct for methodological variation.

### Electron microscopy

Samples for electron microscopy were routinely collected along the experiments and n cells were harvested. ells were then placed in an aluminum disc with a depression of 100 μm and outer diameter of 3 mm (Engineering Office M. Wohlwend GmbH), then covered with a matching flat disc. The sandwiched sample was high-pressure frozen using an EM ICE high pressure-freezing device (Leica Microsystems, GmbH, Germany). Frozen samples were dehydrated in a temperature-controlled AFS2 Freeze substitution device (Leica Microsystems). Substitution was performed in dry acetone containing 1% glutaraldehyde, 1% osmium tetroxide and 0.1% Uranyl Acetate at − 90 °C for 64 h. The temperature was gradually increased to −20 °C (2.9 °C/h) and then raised to 4° C (12 °C/h). The samples were washed five times with acetone and infiltrated for 4 days at room temperature in a series of increasing concentrations of Epon in acetone. After polymerization at 60 °C for 48 h, ultrathin sections (90 nm) were obtained using an EMUC7 ultramicrotome (Leica microsystems) and were mounted on formvar coated 200 mesh nickel grids. Sections were stained with Reynolds lead citrate and examined using a Thermo Fisher Scientific Tecnai T12 transmission electron microscope operating at 120 kV. Digital electron micrographs were acquired using a bottom mounted TVIPS TemCam-XF416 4k x 4k CMOS camera.

## Acknowledgments

We thank Philipp Keller, Julia-Eva Fortmueller, Filipe Natalio, Kang Zhou, Iddo Pinkas, Alla Falkovich, Hortense Viefaure and Pauline Ejsmont for helping with the experiments and Samuel Lovat, Ari Satanowski, Kang Zhou for reading and commenting on the text. Electron microscopy studies were conducted at the Irving and Cherna Moskowitz Center for Nano and Bio-Nano Imaging at the Weizmann Institute of Science. We are grateful to the late Arren Bar-Even for providing the formatotrophic *E. coli* strain.

## Contributions

A.G., A.B.L., E.N., R.M., R.B.-N., E.M and D.F. participated in the project conceptualization and experimental design. D.F., R.B.-N., E.M., G.J., N.D., and G.F. performed the experiments. R.F. performed data visualization. E.N., and R.M. were involved in planning and supervised the work. D.F. wrote the manuscript with input from all authors. All authors commented on the manuscript.

## Notes

### Competing Interest Statement

The authors have declared no competing interest.

### Summary of Updates

Figures 1 and 2 revised, Author contributions updated, Supplemental files updated

