## Supplementary figures and images for "Demonstration of Bioplastic Production from CO_2_ and Formate using the Reductive Glycine Pathway in *E. coli*"

### Supplemental figure 1

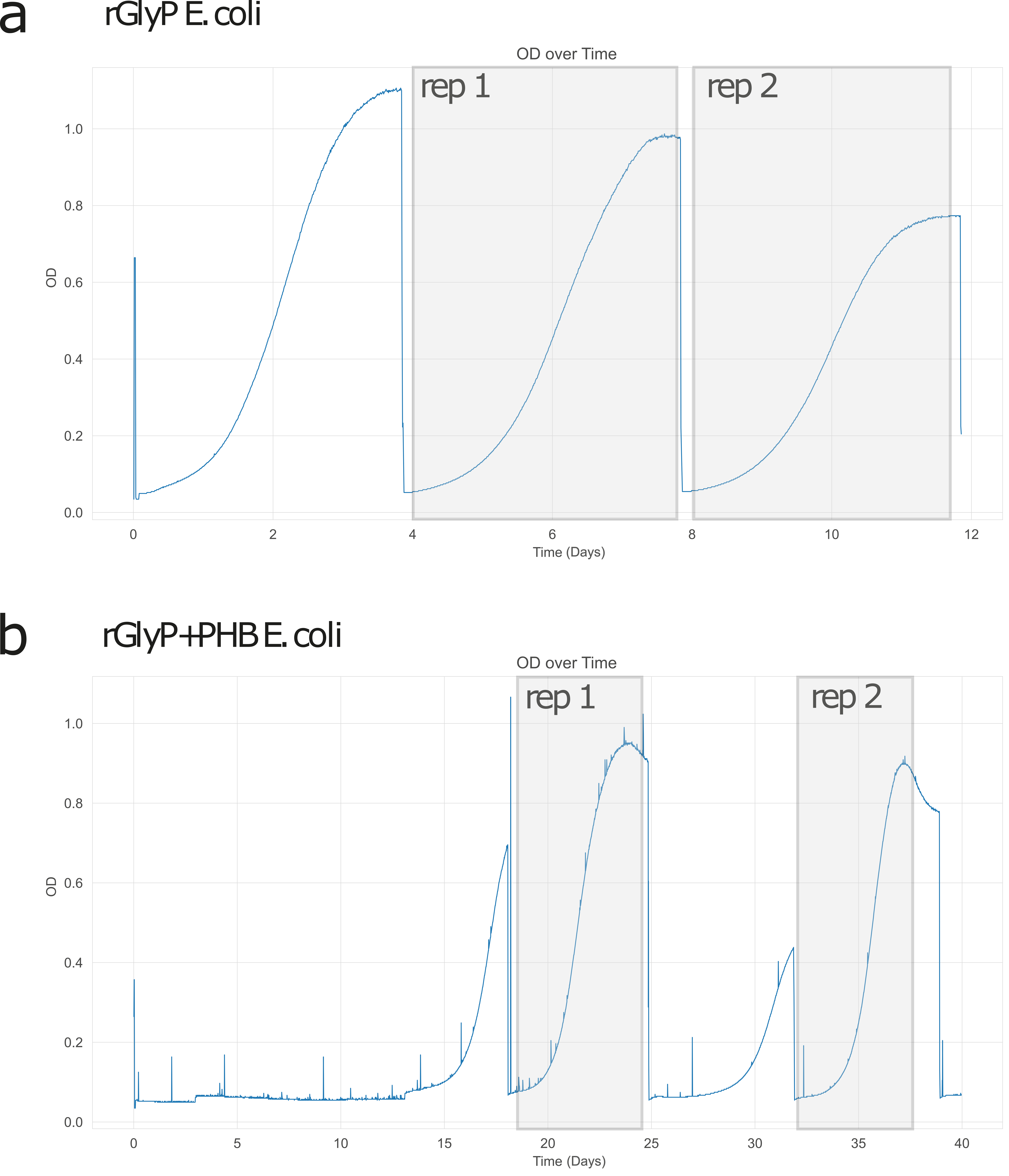

### Supplemental figure 2

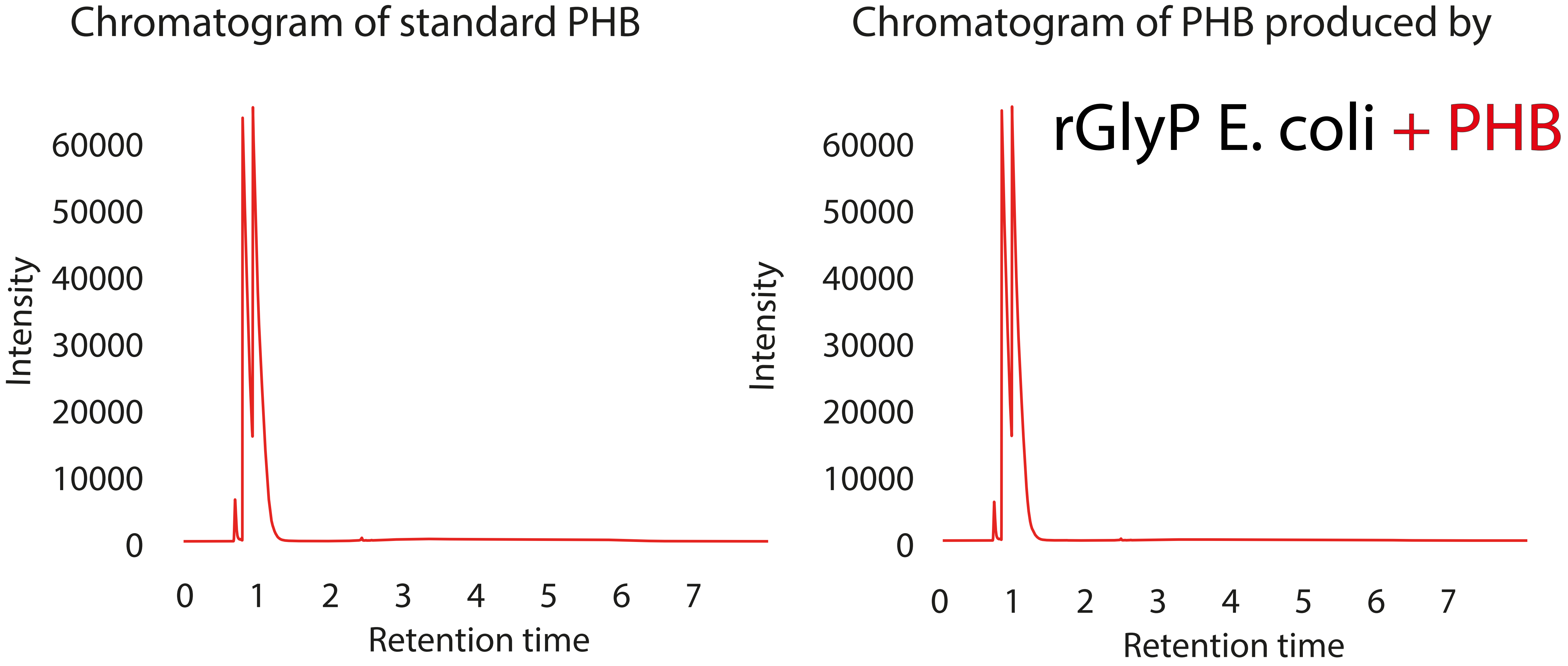
